# Structural basis for prodrug recognition by the SLC15 family of proton coupled peptide transporters

**DOI:** 10.1101/454116

**Authors:** Gurdeep S. Minhas, Simon Newstead

## Abstract

A major challenge in drug development is the optimisation of intestinal absorption and cellular uptake. A successful strategy has been to develop prodrug molecules, which hijack solute carrier (SLC) transporters for active transport into the body. The proton coupled oligopeptide transporters, PepT1 and PepT2, have been successfully targeted using this approach. Peptide transporters display a remarkable capacity to recognise a diverse library of di‐ and tri-peptides, making them extremely promiscuous and major contributors to the pharmacokinetic profile of several important drug classes, including beta-lactam antibiotics, anti-viral and antineoplastic agents. Of particular interest has been their ability to recognise amino acid and peptide-based prodrug molecules, thereby providing a rational approach to improving drug transport into the body. However, the structural basis for prodrug recognition has remained elusive. Here we present crystal structures of a prokaryotic homologue of the mammalian transporters in complex with the antiviral prodrug valacyclovir and the peptide based photodynamic therapy agent, 5-aminolevulinic acid. The valacyclovir structure reveals that prodrug recognition is mediated through both the amino acid scaffold and the ester bond, which is commonly used to link drug molecules to the carrier’s physiological ligand, whereas 5-aminolevulinic acid makes far fewer interactions compared to physiological peptides. These structures provide a unique insight into how peptide transporters interact with xenobiotic molecules and provide a template for further prodrug development.

## Introduction

Solute carrier (SLC) transporters are increasingly being recognised as important determinants of drug efficacy in clinical trials and as important therapeutic targets (1, 2). Poor oral bioavailability is one of the leading causes of compound failure in preclinical and clinical drug development and a major challenge for the pharmaceutical and biotechnology industries (3). A successful approach to address this challenge has been the development of prodrugs that target the intestinal peptide transporter, PepT1 (SLC15A1) (4) (*SI Figure 1*). Prodrugs are bioreversible derivatives of drug molecules that undergo an enzymatic or chemical transformation *in vivo* to release the active parent drug (5). Over the past 10 years significant effort has been made in the design of novel prodrug molecules with improved pharmacokinetic profiles (6). However, targeting specific SLC transporters for carrier mediated uptake is still a major challenge. PepT1 exhibits a remarkably promiscuous binding site and is known to transport many different drug molecules. These include but are not limited to, angiotensin converting enzyme inhibitors, beta-lactam antibiotics, an *N*-methyl-D-aspartate receptor antagonist PD-15874 and 5-aminolevulinic acid, an endogenous non-protein amino acid currently being evaluated as a photodynamic therapeutic agent for the treatment of bladder cancer and Esophageal carcinoma (7-9). Whilst PepT1 is the first peptide transporter encountered by drugs following oral dosage, a second peptide transporter, PepT2 (SLC15A2), functions to selectively reuptake peptides in the nephron and also functions in peptide transport across the blood brain barrier (10). As such, prodrugs targeting both PepT1 and PepT2 show favourable absorption and retention profiles in animal models of drug disposition and are being actively pursued as valid targets for improving pharmacokinetic profiles (11-13).

A major breakthrough in carrier mediated prodrug development was the introduction of the antiviral valacyclovir, marketed under the trade names Valtrex and Zelitrex. Valacyclovir (Cambridge Chemical Database id: TXC) is a prodrug derivative of the antiviral agent acyclovir, which is used in the treatment of disease caused by herpes viruses (including herpes zoster, HSV-1 and −2) as well as in prophylaxis against acquisition of infection and in suppression of latent disease (14). The oral bioavailability of valacyclovir improved to > 50 % for the prodrug derivative valacyclovir compared to 15 % for the parent drug acyclovir, which was attributed to its recognition and transport by PepT1 (15-18). Although the extreme promiscuity displayed by PepT1 has made it a major focus of prodrug strategies (19-21), the structural basis for prodrug recognition is still enigmatic. Lack of structural information on how prodrugs interact with the transporter is hampering efforts to design accurate pharmacophore models for, amongst other developments, computer aided drug design (22, 23).

To date bacterial peptide transporters have proven to be valid model systems with which to understand the molecular basis of peptide recognition within the human PepT1 and PepT2 transporters (24-26). PepT1 and PepT2 belong to the much larger POT or PTR family of proton coupled oligopeptide transporters, with homologues found in all domains of life except the archaea (27, 28). POT family transporters belong to the Major Facilitator Superfamily (MFS) of secondary active transporters, and use the proton electrochemical gradient to drive the concentrative uptake of di‐ and tri-peptides into the cell (29). Although crystal structures of bacterial POT family transporters have revealed that peptides can be accommodated in distinct binding positions (30, 31) and transported with variable proton stoichiometries (32), the rules governing how specific functional groups on peptides are recognised remain obscure.

To address this question and understand how peptide-based prodrugs and non-proteinogenic drug molecules are recognised and transported via the SLC15 family, we determined the crystal structure of a bacterial POT family member in complex with both valacyclovir and 5-aminolevulinic acid at 3.1Å and 2.5 Å resolution respectively. Combined with the previous peptide bound structures, we present a pharmacophore model for prodrug recognition, which facilitates a structure-based route to further drug development targeted at the SLC15 family.

## Results and Discussion

### Structure of valacyclovir complex.

Recently we discovered a prokaryotic homologue of PepT1 from the bacterium *Staphylococcus hominis*, PepT_Sh_, which was able to transport a natural thioalcohol conjugated peptide, exhibiting structural characteristics similar to prodrug molecules (*SI Figure 2*) (33). We therefore reasoned that PepT_Sh_ may also be able to recognise and transport valacyclovir. Using a competition assay we determined that valacyclovir was able to compete for di-peptide binding in PepT_Sh_ with an IC_50_ value of 7.4 mM (Figure 1A). Although higher than equivalent IC_50_ values obtained for natural peptides, which were 72.2 µM for di-Alanine (AlaAla) and 23.7 µM for tri-Alanine (AlalAlaAla), similar K_M_ values have been reported for valacyclovir uptake in mouse and human PepT1 (34, 35). We were further able to measure both valacyclovir and 5-aminolevulinic acid transport directly using a pyranine based transport assay that measures proton movement (32) (Figure 1B & C), supporting the use of PepT_Sh_ as a model for understanding prodrug transport in the mammalian proteins.

**Figure 1.**
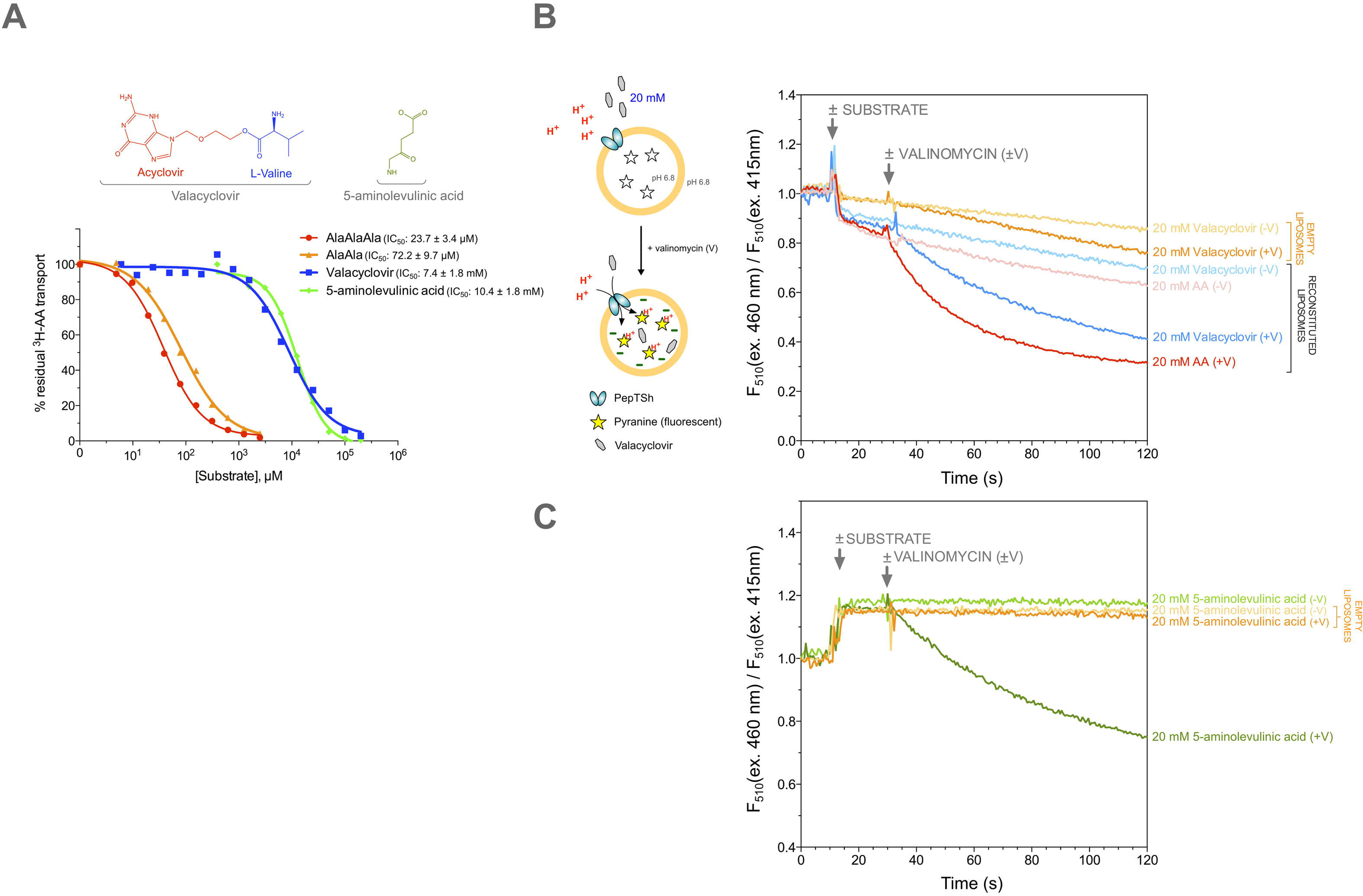
Valacyclovir and 5-aminolevulinic recognition by PepT_Sh_. (A) Chemical structures of valacyclovir and 5-aminolevulinic acid. The L-valine scaffold is shown in blue and active drug molecule in red. IC_50_ competition curves for dipeptide (AlaAla), tripeptide (AlaAlaAla), valacyclovir and 5-aminolevulinic acid. (B) Measurement of proton coupled uptake of valacyclovir using a pH reporter assay. PepT_Sh_ was reconstituted into liposomes loaded with pyranine and a high concentration of potassium ions. The external solution contains either peptide, AlaAla (red) or valacyclovir (dark blue) and a low potassium concentration. On addition of valinomycin (+V) the membrane becomes highly permeable to potassium, generating a hyperpolarised membrane potential (negative inside), which drives transport, protonating the pyranine dye and reducing emission at 510nm following excitation at 460nm. (C) Measurement of proton coupled uptake of 5-aminolevulinic acid using a pH reporter assay, experimental set up is as shown in (B).

Following extensive screening, a crystal structure of PepT_Sh_ in complex with valacyclovir was subsequently determined using the *in meso* crystallisation method (36) and refined to a final resolution of 3.1 Å (*SI Table1*). PepT_Sh_ adopts an almost identical inward open conformation to that obtained in our previous study (33), with a root mean square deviation (r.m.s.d) of 0.499 Å for 480 C_β_ atoms (*Figure 2A*). The valacyclovir drug molecule was clearly observed sitting in the central peptide binding site, coordinating a single water molecule (*Figure 2B*). The valine end of valacyclovir orientates towards the extracellular gate (TMs 1,2 & 7,8), which is closed, and the nucleoside part of the drug orientates towards the intracellular gate (TMs 4,5 & 10, 11), which is open.

**Figure 2.**
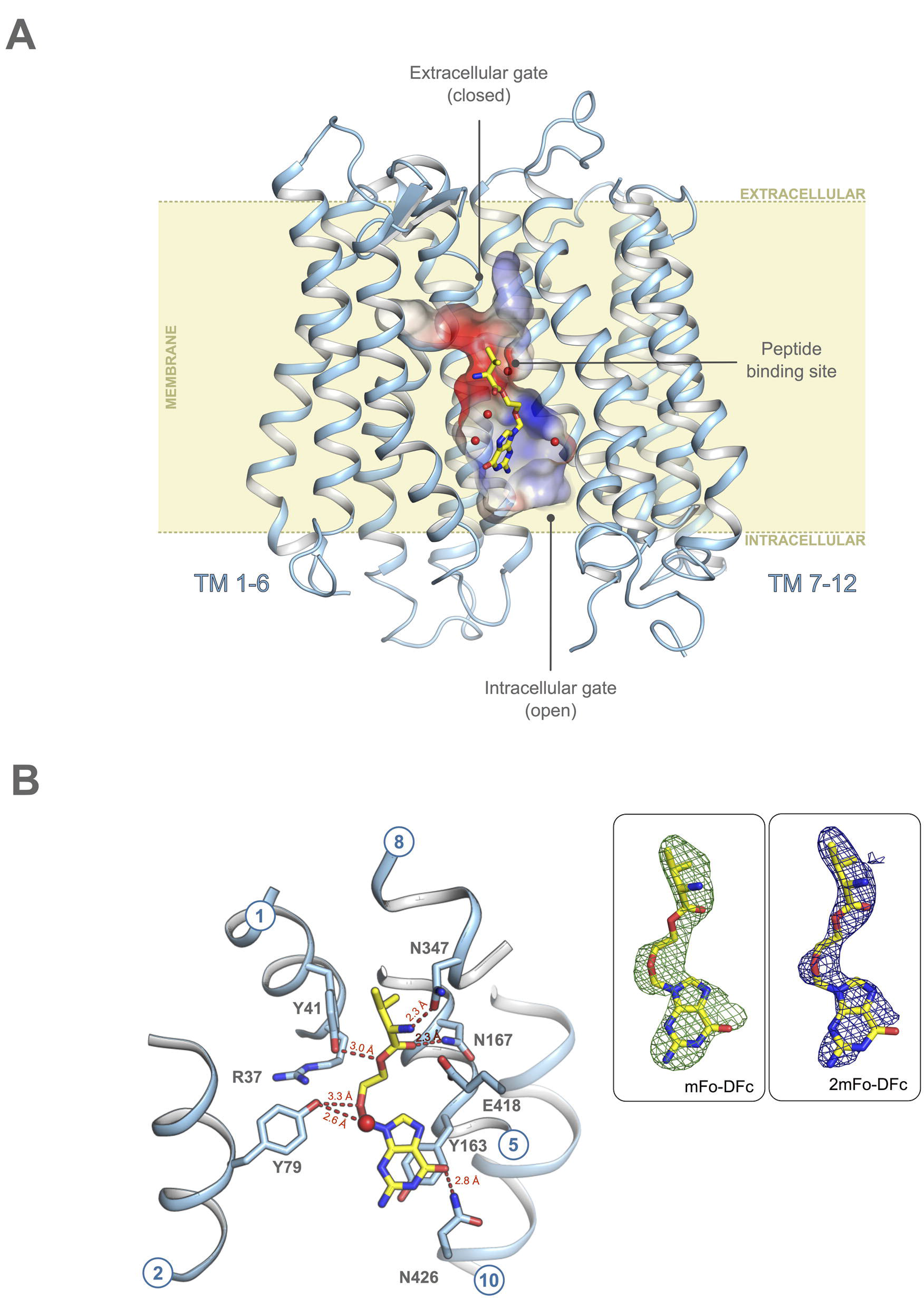
Crystal structure of PepT_Sh_ in complex with valacyclovir. (A) PepT_Sh_ represented as light blue helices in the plane of the membrane. The binding cavity surface is coloured according to the localised electrostatic potential. Yellow sticks represent the bound prodrug, valacyclovir. Waters are shown as red spheres. (B) Residues that interact with valacyclovir are shown as blue sticks. Hydrogen bond are shown as red dashes and distances labelled. Inset: Experimental *mFo-DFc* difference electron density (green mesh) observed for valacylcovir bound to PepT^Sh^ contoured at 3 σ. Final refined 2mFo-Fc electron density map, blue mesh, contoured at 1 σ, (PDB:6GZ9)

Valacyclovir makes a number of specific interactions to side chains that are strictly conserved within the wider SLC15 family (Figure 3 and *SI Figure3*). The amino terminus interacts with a conserved glutamate, E418 (E595 – human PepT1 numbering) on TM10 and through a hydrogen bond to an asparagine on TM8, N347 (N329). These residues are essential for binding and transport of peptides in both mammalian and bacterial POT family transporters (37, 38). The carbonyl group makes a hydrogen bond to a conserved asparagine, N167 (N171) on TM5, and the ester bond, which links the L-valine to the acyclovir, interacts with tyrosine Y41 (Y40) on TM1. The acyclovir ether group interacts with another conserved tyrosine, Y79 (Y64) on TM2 and the nucleoside portion of the drug makes a distinctive pi-pi stacking interaction with tyrosine Y163 (Y167), also on TM5. Equivalent tyrosine residues play important roles in determining the affinity and transport of peptides in the mammalian proteins (39). The only non-conserved interaction is made between the nucleoside hydroxyl and N426 on TM10, which in PepT1 is a leucine (L630). All these helices are part of the gating architecture within the POT family, which must couple ligand binding to proton movement during transport (40). Of note are the interactions made by the amino terminus of the L-valine scaffold to E418 and N347 and the carbonyl group to N167, as these closely mimic those observed previously in complexes of a related POT family transporter, PepT_St_, to natural peptides (26, 30, 41) and the mammalian homologue PepT1 (42).

**Figure 3.**
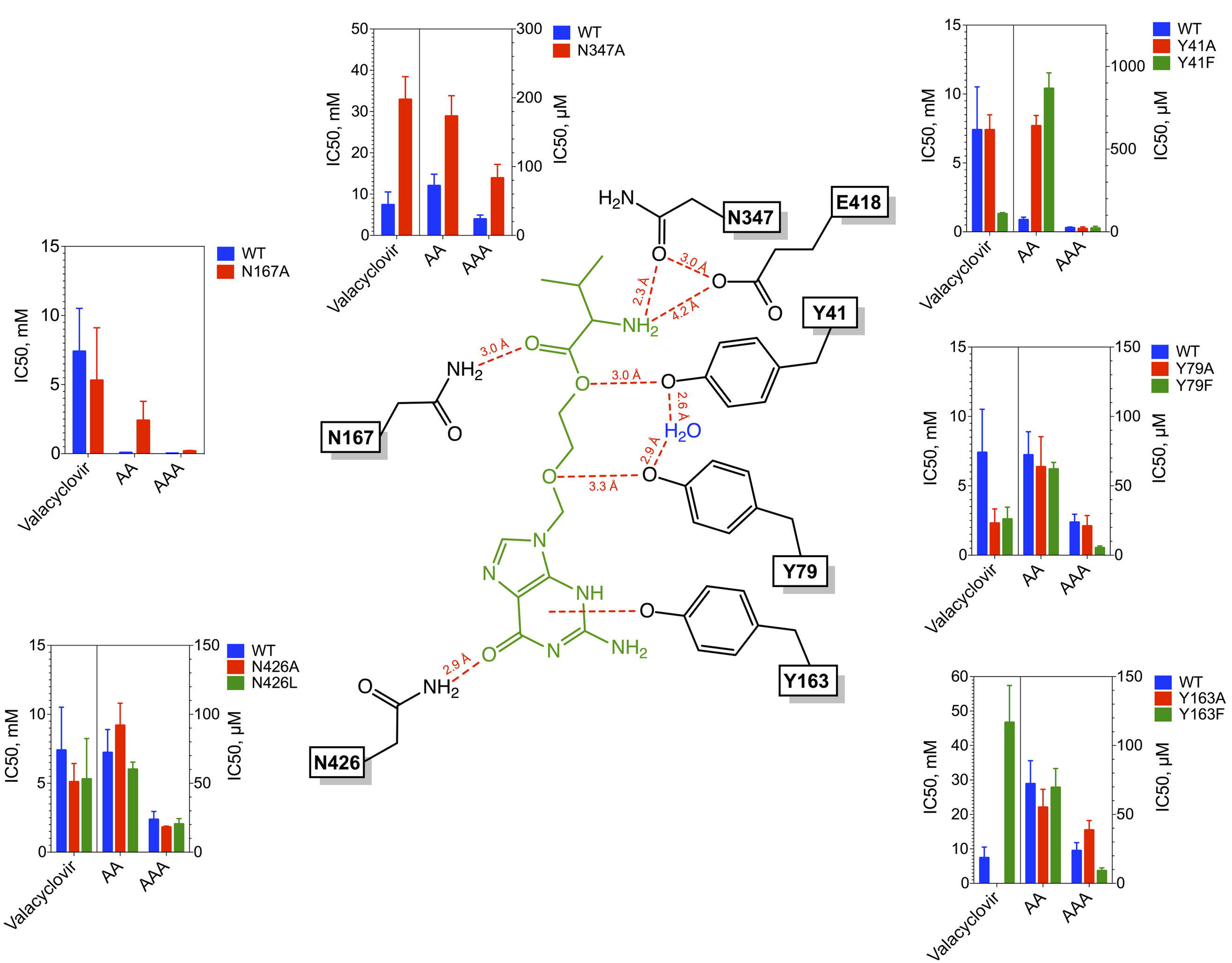
Functional analysis of PepT_Sh_ binding site variants. Schematic of valacyclovir (green) interacting with PepT_Sh_ (black). The contribution of each interacting residue was tested by mutagenesis. The IC_50_ for each mutant was measured for three substrates: AlaAla, AlaAlaAla and valacyclovir and plotted as a bar graph compared against WT (blue bars).

### Functional analysis of binding site residues contributing to valacyclovir recognition.

To further understand the functional importance of these interactions we undertook a detailed functional analysis of the binding site residues (*Figure 3, SI Table 2 and SI Figure 4*). Previous studies have shown that interactions to the amino terminus of natural peptides are well conserved within the POT family, as are the interactions to the carbonyl oxygen (29). We tested the importance of the N-terminal interactions in PepT_Sh_ by measuring uptake of either N‐ and C-terminally blocked peptides (*SI Figure 5*). Only the free N-terminal peptide was recognised, consistent with previous reports for the mammalian transporter (38). Further underlining the importance of the N-terminal interactions, a reduction in binding affinity for the N347A variant was also observed (*Figure 3 & SI Table 2*). Mutations of E418 on the other hand resulted in an inactive transporter (*SI Figure 4*), consistent with previous results showing the essential role of the TM10 glutamate in controlling the intracellular gate in response to peptide and proton binding (37, 41).

Prodrugs often contain ester groups, as these confer labile linkages between the active drug molecule and the scaffold, which are easily cleaved once the prodrug has been transported across the membrane (43). An important question is how these functional groups are accommodated within the peptide transporter binding site. The ester linkage interacts through a hydrogen bond with Y41, which plays an important role in peptide recognition and forms part of the conserved E^33^xxERFxYY sequence motif on TM1 (37, 44). However, removing either Y41 or N167, which interacts with the carbonyl group close to the ester linkage, had little effect on the affinity for valacyclovir, whereas a conservative phenylalanine substitution at Y41 resulted in a decrease of the IC_50_ from 7.4 to 1.3mM. Interestingly, we observe a similar decrease in IC_50_ values for valacyclovir to 2.2mM for the Y79F variant, which interacts with the ether linkage. These results suggest that while specific interactions to the ester and ether groups in valacyclovir are made, these are not required for prodrug recognition and are merely accommodated within the binding site.

Interestingly however we did observe substantial differences in the IC_50_ values between di‐ and tri-peptides in several of the variants tested (Figure 3 & SI Table 2). In particular Y41 appears to play a more important role in di-peptide recognition, as replacement with phenylalanine resulted in an increase in IC_50_ from 72.2 µM to 866 µM whilst no effect was observed for tri-alanine. A similar result M M, was obtained for the Y41A variant. In the Y79F variant however, we observed the opposite trend, with limited effect on di-alanine transport but a positive effect on tri-alanine recognition, with a reduction in IC_50_ from 23.7 µM o 5.3 µM he results from the N167A variant also showed a differential effect on di‐ and tri-alanine, with the latter being impacted to a far greater degree. These results lend further support to our hypothesis that peptides are accommodated in different positions within the binding site (29) and that this mechanism is shared more widely within the POT family. As we discuss below, a general mechanism for accommodating peptides in different orientations is also supported by the comparison of the crystal structures of valacyclovir and 5-aminolevulinic acid with previous peptide bound complexes.

Peptide transporters accommodate the chemical diversity of side chain groups within specificity pockets, which contain several conserved tyrosine and polar side chains (29). Unexpectedly the non-protein purine ring of valacyclovir does not occupy one of these pockets, as previously suggested (45). However, a favourable interaction is observed through a conserved tyrosine, Y163, via pi-pi stacking and with a non-conserved asparagine, N426, through the carbonyl group of valacyclovir. Y163 (Y167) forms part of the PTR2_2 signature motif in the SLC15 family and plays an important role in peptide recognition in human PepT1 (27, 28). With reference to valacyclovir we can extend this function to reveal a key role for tyrosine 163 in accommodating the purine ring. Removal of Y163 abrogated valacyclovir recognition, underscoring its importance. The more conservative phenylalanine substitution also had a negative effect on valacyclovir recognition, increasing the IC_50_ from 7.4 mM to 40 mM. This highlights the importance of the pi-cation interaction between the purine ring and the phenolic hydroxyl group. Removal of N426 again resulted in an improved affinity for valacyclovir, in line with previous results for Y41 and Y79, whereas replacing the side chain with leucine, which is found in the human transporter had a negligible effect. It is likely the interaction with N426 is specific for PepT_Sh_ and does not occur in the human transporter.

### Structure of 5-aminolevulinic acid complex

We successfully captured PepT_Sh_ in complex with 5-aminolevulinic acid and determined two structures at 2.5 and 2.8 Å resolution respectively (*Figure 4A and SI Table 1*). 5-aminolevulinic acid is an endogenous non-protein amino acid that forms the first part of the porphyrin synthesis pathway in mammals (46) and is used in the clinic for the photo dynamic detection and treatment of cancer (47). The IC_50_ value for 5-aminolevulinic acid in PepT_Sh_ is 10.4 mM, which is similar to that obtained for valacyclovir of 7.4 mM (*Figure 1A*), and within the same range observed in PepT1, of 2.1 mM (48). PepT_Sh_ adopts an almost identical inward open state observed previously, with an r.m.s.d of 0.305 Å over 480 C_α_ atoms when compared with the valacyclovir structure. However, similar to the valacyclovir structure the B-factors and quality of electron density around the cytoplasmic facing regions of TM10 and 11 clearly indicate increased flexibility in this part of the transporter structure (SI Figure 6). This is consistent with previous results obtained for other POT family transporters and lend further support to the C-terminal bundle being more dynamic in this family of MFS proteins (37, 49).

**Figure 4.**
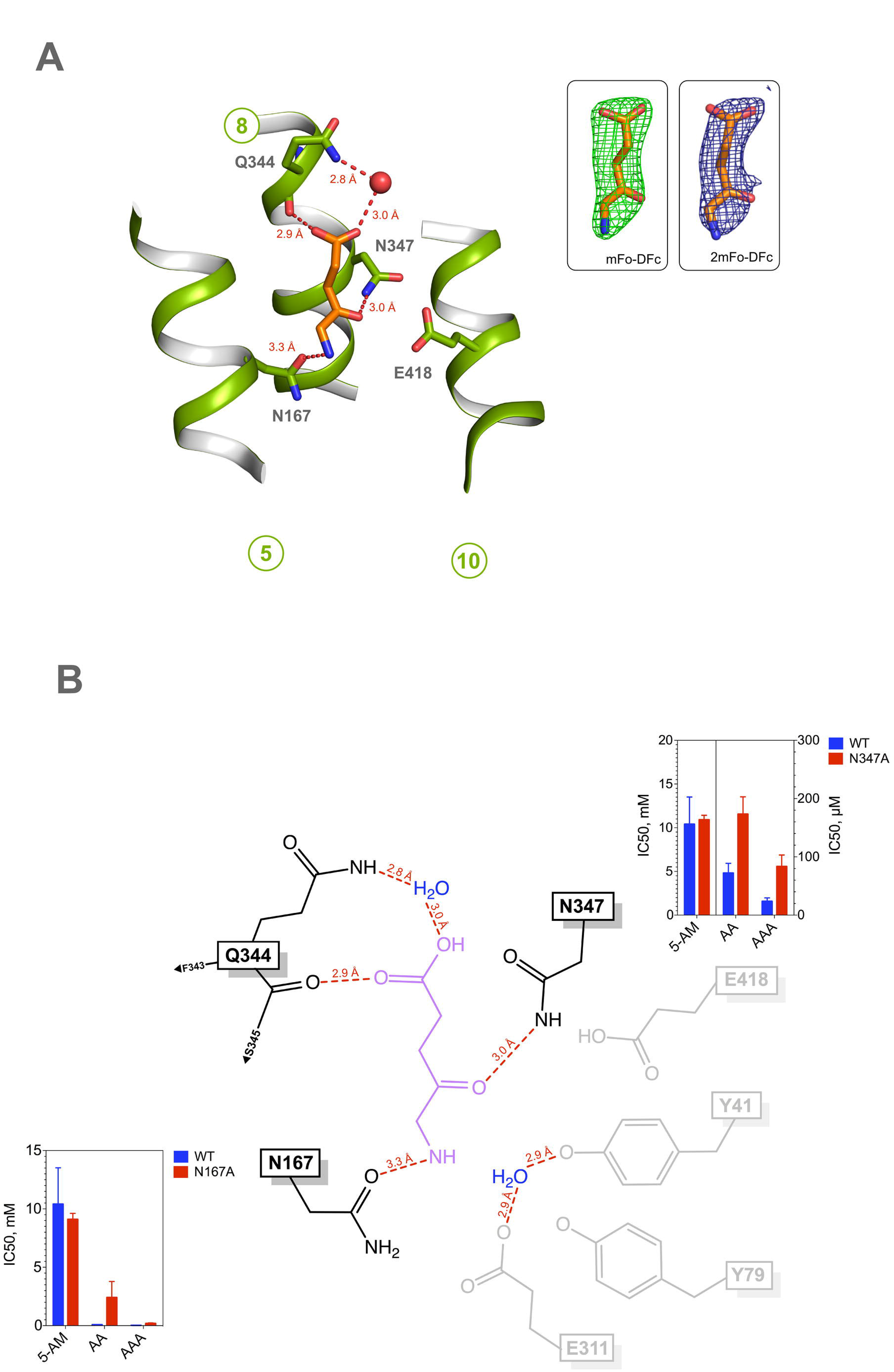
Crystal structure of PepT_Sh_ in complex with 5-aminolevulinic acid. (A) 5-aminolevulinic acid (orange sticks) bound to PepT_Sh_ (green) (PDB:6HZP). Hydrogen bond interactions are shown as red dashes with distances indicated. Water molecules are shown as red spheres. Key residues involved in binding substrate are shown as green sticks. Inset: Experimental *mFo-DFc* difference electron density (green mesh) observed for 5-aminolevulinic acid, contoured at 3 σ. Final refined 2mFo-Fc electron density map, blue mesh, contoured at 1 σ. (B) Schematic of 5-aminolevulinic acid (purple) interacting with PepT_Sh_ (black). Nearby residues are indicated in grey, but are not interacting with the ligand. IC_50_ values for the two variants tested are shown as bar charts and compared to WT values.

The 5-aminolevulinic acid molecule sits in a similar position as the L-valine scaffold in valacyclovir, adopting a vertical orientation (*SI Figure 7*). However, there are notable differences between how the two molecules interact with the binding site. The C-terminal carboxyl group of 5-aminolevulinic acid faces towards the extracellular gate, making interactions with the backbone carbonyl group of Gln344 (Gln326) and via a water mediated bridge to the side chain of this same residue. Gln344 forms part of a highly conserved sequence motif at the extracellular part of TM8 in the mammalian peptide transporters (PDQMQ), but only the glutamine is observed in PepT_Sh_ (*SI Figure 8*). The interaction with a backbone carbonyl group and water molecules is similar to the tri-Alanine structure observed in PepT_St_ (30) (PDB: 4D2D), suggesting the vertical mode of binding is associated with lower affinity compared to the more horizontal configuration.

5-aminolevulinic acid does not contain a peptide bond, having a ketomethylene group instead. This sits in close proximity to E418, in a similar position to the amino terminal group of the L-Valine scaffold in valacyclovir. The amino terminal group in contrast sits in a similar position to the carboxyl group of valacyclovir, interacting with N167. Unexpectedly the IC_50_ value for the N167A variant was unchanged (*Figure 4B*) suggesting this interaction is not essential for binding. It is possible that 5-aminolevulinic acid can form compensatory interactions in the N167A variant or that in the vertical position, the carboxyl group provides compensatory interactions.

Finally, during refinement it became clear that an unusual interaction could be observed within the binding site, wherein an arginine on TM1 (R37) and lysine on TM 4 (K137) interact through a shared hydrogen bond (*SI Figure 9*). Arginine 37 forms part of the conserved E^33^xxERFxYY motif on TM1 (*SI Figure 2*), which was previously identified as playing an important role in the proton coupling mechanism in the POT family (37). Indeed, we previously postulated a role for the conserved arginine in regulating the pKa of the TM4 (25). However, the current structure provides the first experimental evidence of a direct interaction between these two side chains. However, we could not discern any significant influence of 5-aminolevulinic acid on the binding site that would cause this interaction to occur, or indeed, of the valacyclovir to break this interaction. Further analysis will be needed to follow up this observation.

## Discussion

Previously determined crystal structures of bacterial peptide transporters in complex with di‐ and tri-peptides have revealed key features of how these ligands are recognised within the POT/SLC15 family (26, 30, 31, 41). To identify commonalities between peptide and prodrug recognition and develop a pharmacophore model for prodrug binding, it is instructive to compare the valacyclovir binding mode with previous peptide based complex structures. Superposition of the di-peptide bound crystal structures of the SLC15 transporter from *Streptococcus thermophilus*, PepT_St_ (30, 31) reveal several noticeable commonalities in the binding position of the prodrug and natural peptides (*SI Figure 10*). The L-valine part of valacyclovir makes very similar interactions to the di-peptides L-Ala-L-Phe and L-Ala-L-Gln, whilst adopting the more vertical orientation observed for the L-Ala-L-Ala-L-Ala tri-peptide. The interactions to the amino terminus are well conserved, as is the hydrogen bond to the carbonyl oxygen through N167. Of particular note is the ester linkage part of the valacyclovir prodrug, which closely matches the position of the peptide bond in the di-peptide structures. Indeed, the equivalent tyrosine to Y41 in PepT_St_, which we observe being made to the ester bond in valacyclovir, makes a similar interaction to the amide nitrogen in the peptide bond in the di-peptide complex. It was known previously that although peptide bonds are not strictly required for recognition in PepT1, the presence of a carbonyl group within close proximity to the amino terminus is an important feature of high affinity ligands (12). It is interesting to note that whilst neither valacyclovir nor 5-aminolevulinic acid have a peptide bond, they still present hydrogen bond acceptors or donors to both N167 and E418, satisfying this requirement.

Valacyclovir does not contain a terminal carboxy group, which in the di-peptide ligand can be seen making favourable electrostatic interactions to two conserved positively charged side chains in the N-terminal bundle. In valacyclovir we observe the ether bond occupying a similar spatial position as the peptide carboxyl group. It is likely the close placement to arginine 37 (R27) facilitates accommodation of the ether group, however, another interaction to a conserved tyrosine, Y79, is observed.

Of particular interest was the observation that 5-aminolevulinic acid adopted a vertical orientation as opposed to the horizontal one adopted by dipeptides in PepT_St_ (30, 31). It is unclear whether this is due to the presence of the ketomethylene group replacing the peptide bond or the absence of side chains that would be accommodated within the specificity pockets found in the binding sites. A similar vertical orientation was observed for a bound thioalcohol peptide, Cys-Gly-3M3SH, in our previous structure of PepT_Sh_ (33) and a common set of interactions between these three ligands can be discerned, being made to Y41 and N167 (SI Figure 11), which are strictly conserved throughput the SLC15 family (SI Figure 2). To a lesser extent we also observe interactions to N347, the backbone carbonyl of Q344 and several ordered water molecules, which also play an important role in proton movement within peptide transporters (50). However, recognition of the unusual Cys-Gly-3M3SH peptide appears to be largely driven through accommodation of the thioalcohol group in an extended hydrophobic pocket, which seems to be an evolutionary adaption unique to PepT_Sh_ and its environmental niche as a skin commensal (33).

Our understanding of which scaffolds make optimal candidates for prodrug development is still evolving (6, 22). However, the comparison of the three ligands in PepT_Sh_, and their analysis with respect to peptides bound in PepT_St_, suggests that common points of interaction between the transporter and different ligands exist, which may present a novel route for prodrug scaffold design.

### Pharmacophore model for valacylcovir binding

Taken together these results enable us to propose a structure-based pharmacophore model for valacyclovir binding to peptide transporters (*Figure 5*). We are more circumspect regarding a model for 5-aminolevulinic acid however, as our current mutagenesis data provides less information for this drug. Nevertheless, the structural comparison of valacyclovir with previous di-peptide co-crystal structures suggests a significant contribution to recognition is made through the amino terminus of the scaffold L-Valine, which interacts with N347 and E418. We observe a similar pattern of interactions with the carbonyl group of both the ester bond and the peptide bond to N167. A surprising finding was that while the binding mode of valacyclovir closely replicated that of physiological peptides, the contributions of the interactions to the affinity were noticeably different. Most surprising was that removing the interactions to conserved side chains N167, Y41 and Y79 increased the affinity for valacyclovir, whilst having a generally negative effect on the transport of di‐ or tri-alanine. Importantly, a similar increase in the IC_50_ values for valacylcovir were observed in equivalent variants in the related bacterial peptide transporter from *Shewanella oneidensis*, PepT_So_ (51) (SI Table 3), lending support to a common mechanism of prodrug recognition within the SLC15 family. Valacyclovir is however much larger than a tri-peptide, making the physical constraints of accommodating this molecule more severe. Indeed, the drug appears to occupy a position halfway between the horizontal and vertical poses previously observed in PepT_St_ (30, 31).

**Figure 5.**
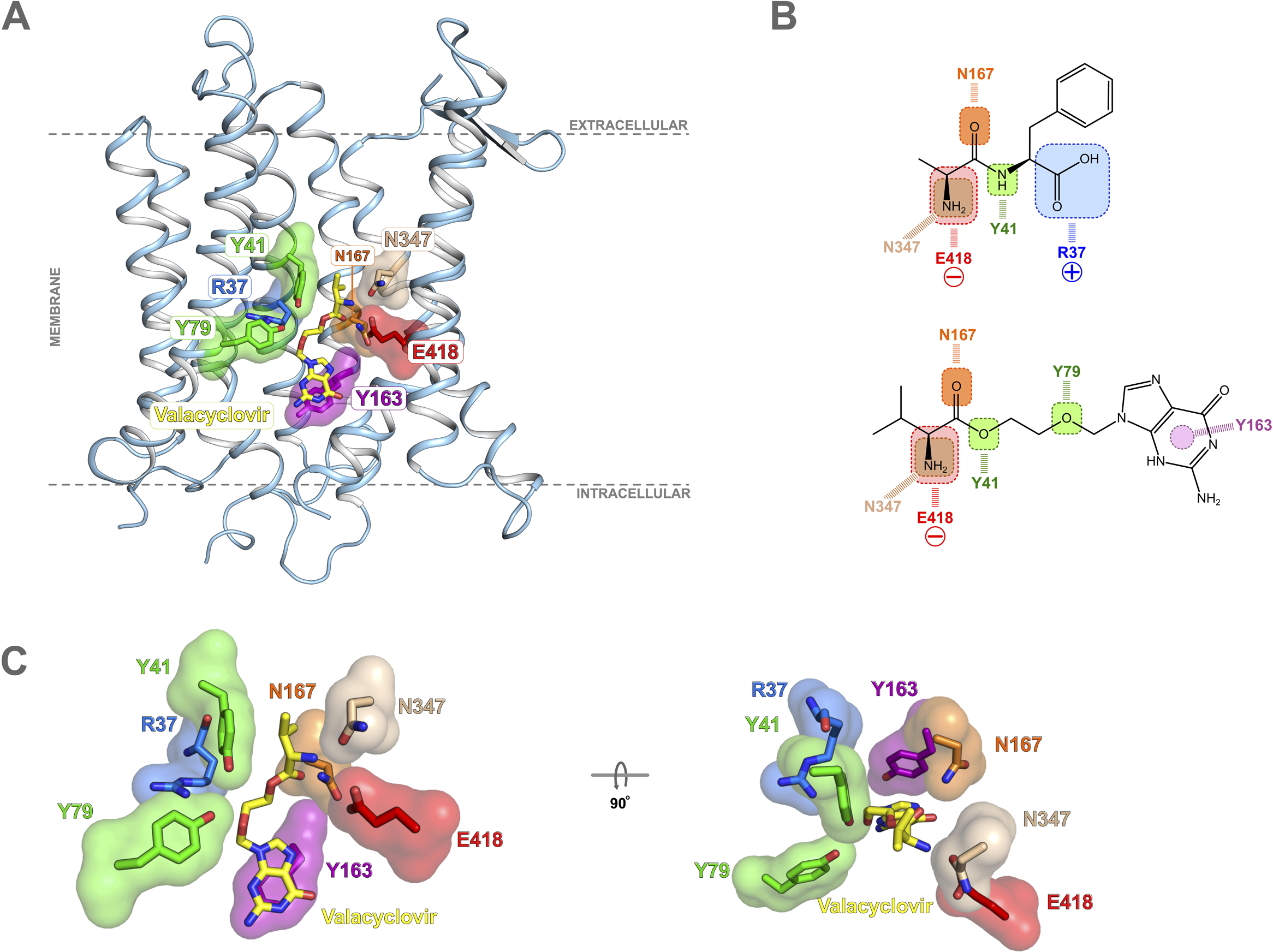
Pharmacophore model for valacyclovir binding to POT family transporters. Key interaction sites observed in PepTSh are shown for valacyclovir in the context of the full transporter structure. (B) Proposed pharmacophore model indicating the role of conserved SLC15 family binding site residues in recognising either peptides (AlaPhe) (PDB: 4D2C) or prodrug (valacyclovir). (C) Close up view into the PepTSh binding site accommodating valacyclovir highlighting interactions shown in (A).

A noticeable difference between the valacyclovir and peptide complexes was also observed in the region corresponding to the free carboxyl group. The carboxy terminal group in di-peptides interacts electrostatically with two conserved positively charged side chains in the N-terminal bundle (30). The dipole made between these and the conserved glutamate, E418 on TM10 (E595) helps to orientate smaller peptides in the binding site. The absence of a carboxyl group in valacyclovir results in no interactions being made to the positive cluster in the N-terminal bundle. However, a free carboxyl group is not required for peptide recognition in mammalian PepT1 (38) or PepT_Sh_ (*SI Figure 5*), explaining why this interaction is not strictly required. A further important observation was the role played by Y163 in accommodating the non-peptidometic purine ring within the binding site. This result supports a broader role for tyrosine side chains in contributing to the drug binding within the human peptide transporters. Additionally, the structural comparison with valacyclovir further develops our previous hypothesis that the promiscuity observed in peptide transporters stems in part from their ability to accommodate ligands in different orientations in the binding site (29). This ‘multi-mode binding model’ for ligand recognition would also explain why prodrug molecules can be accommodated through interactions with the amino terminal group on the scaffold and steric complementarity between the ester linkage and peptide backbone.

### Concluding remarks.

The development of prodrugs has progressed with the aim of improving drug pharmacokinetics by overcoming various barriers that reduce clinical efficacy, such as poor oral bioavailaibility or cellular toxicity due to adverse drug-drug interactions (1). Carrier mediated prodrug design has been successfully employed to overcome these challenges (6), however prodrug design remains difficult owing to the lack of a generally applicable strategy that can be broadly applied. The unusual promiscuity of PepT1 and PepT2 make them ideal targets for prodrug development (7). The current structures and the associated pharmacophore model for valacyclovir and 5-aminolevulinic acid recognition by PepT_Sh_ provides an important advance in our attempts to rationalise and exploit PepT1 and PepT2 more widely in carrier mediated drug transport.

### Methods.

The gene encoding *PepT_Sh_* was cloned into a C-terminal octa-histidine GFP fusion vector (Addgene plasmid 75759) and transformed into Escherichia coli C43(DE3) cells (52). Mutations were introduced using a modified quickchange mutagenesis protocol. An overnight starter culture was used to inoculate 4 L of Terrific Broth (TB). Cells were left to grow at 37°C and aerated by shaking at 250 rpm. Overexpression of the fusion protein was induced when the culture reached an OD_600_ of ~0.6, by adding isopropyl-β-D-thiogalactopyranoside (IPTG) to a final concentration of 0.4 mM. Following induction, the temperature was reduced to 25 °C. Cells were harvested 16 hours later, and resuspended in phosphate buffered saline (PBS) containing deoxyribonuclease I from bovine pancreas (Sigma), and stored at −80°C.

Unless stated, the following steps were carried out at 4 °C. Cells were thawed and disrupted at 32 kpsi (Constant Systems, UK). Cellular debris was removed by centrifugation at 30,000 *xg* for 30 minutes, and membranes were isolated by ultracentrifugation at 234,000 *xg* for 120 minutes. Membranes were resuspended in PBS using a dounce homogeniser, flash frozen, and stored at −80°C.

Thawed membranes were diluted with 1x PBS (10 mL per gram of membrane), 20 mM imidazole (pH 8.0), 150 mM NaCl, and 1 % n-Dodecyl-β-D-maltoside (Glycon) and left to solubilise for 60 minutes, stirring. Non-solubilised membranes were removed by ultracentrifugation at 234,000 *xg* for 60 minutes. Nickel-NTA Superflow resin (ThermoFisher Scientific) was added to the sample, using a ratio of 1 mL resin per gram of membrane, and left to bind, stirring, for approximately 90 minutes. The resin was then packed into a glass econo-column (BioRad, USA) under gravity. The resin was washed with 20 column volumes (CV) of wash buffer (1x PBS, 20 mM imidazole (pH 8.0), 150 mM NaCl and 0.05% DDM). The resin was further washed with 10CV of wash buffer containing 30 mM imidazole.

The bound GFP-PepT_Sh_ fusion protein was eluted with wash buffer containing 250 mM imidazole. The eluent was combined with an equivalent concentration of hexa-histidine tagged TEV protease and dialysed overnight against 20 mM Tris-HCl (pH 7.5), 150 mM NaCl, and 0.03% DDM. The cleaved protein was filtered using a 0.22 μm filter (Millipore, USA) and passed through a HisTrap column (GE healthcare) to remove uncleaved GFP-PepT_Sh_ protein and TEV protease. The flow through was collected and concentrated to 300 μl using a 50 kDa MWCO concentrator (Vivapsin 20, Sartorius AG) and applied to a size exclusion column (Superdex 200 10/30, GE Healthcare) pre-equilibrated with 20 mM Tris-HCl (pH 7.5), 150 mM NaCl, and 0.03% DDM. Fractions containing purified PepT_Sh_ protein were pooled and concentrated to a final concentration of 10-15 mg ml^−1^.

### Crystallisation and Structure Determination.

PepT_Sh_ was incubated with either 40 mM valacyclovir-hydrochloride (Sigma Aldrich) or 5-aminolevulinic acid and left for 4 hours at 4 °C prior to crystallisation. The protein-laden mesophase was prepared by homogenizing monoolein (Sigma) and 10^−1^mg.ml protein solution in a 60:40 ratio by weight using a dual syringe mixing device at 20 °C (53). Crystallisation was carried out at 4 °C in 96-well glass sandwich plates with 50 nL mesophase and 0.8 µM pitant solution. The crystallisation solution consisted of 26-27% (v/v) PEG 200, 220 mM (NH4)_2_HPO_4_, and 110 mM sodium citrate (pH 5.0). Crystals grew within two to three days. Wells were opened using a tungsten-carbide glasscutter and the crystals were harvested using 100 µm micromounts (MiTeGen). Crystals were cryo-cooled directly in liquid nitrogen.

Date were collected at beamline I24 (Diamond Light Source Facility, Oxfordshire, United Kingdom) and ID23eh2 (European Synchrotron Radiation Facility). The data were processed and scaled in XDS (54) and AIMLESS (55).

### Model Building and Refinement.

The structure was phased by molecular replacement in PHASER (56) using the previously resolved PepT_Sh_ crystal structure, PDB: 6EXS. Ligand restraints were generated using Grade Web Server (http://grade.globalphasing.org/). The model was built into electron density using COOT (57) followed by refinement in Phenix (58), and validated using the Molprobity server (59). Figures were prepared using PyMOL (Schrödinger, LLC).

### Fluorescence based transport assays.

Purified PepT_Sh_ was detergent exchanged into n-Decyl-β-D-maltopyranoside (DM) using a size exclusion column (Superdex 200 10/30, GE Healthcare) pre-equilibrated with 20 mM Tris-HCl (pH 7.5), 150 mM NaCl, and 0.3% DM. Lipids (Avanti Polar Lipids) (POPE:POPG in a 3:1 ratio) were mixed with the eluted protein at a ratio of 60:1. The protein:lipid mixture was rapidly diluted with 50 mM KPO_4_ (pH 7.0) buffer. Proteoliposomes were harvested by ultracentrifugation (234,000 *xg*) for 120 minutes. The pellet was resuspended in 50 mM KPO_4_ and dialysed overnight against 3 L of 50 mM KPO_4_ (pH 7.0). Proteoliposomes were recovered and subjected to three rounds of freeze thawing before storage at −80°C.

Proteoliposomes were thawed and harvested by ultracentrifugation at 150,000 *xg*. The resulting pellet was carefully resuspended in 120 mM KCl, 2 mM MgSO_4_, 5 mM HEPES pH 6.8 (INSIDE buffer) and 1 mM pyranine (trisodium 8-hydroxypyrene-1,3,6-trisulfonate) (Sigma). Proteoliposomes then underwent eleven freeze-thaw cycles in liquid nitrogen before the sample was extruded through a 0.4 μm polycarbonate membrane and harvested by ultracentrifugation as previous. Excess pyranine dye was removed with a Microspin G-25 column (GE Healthcare) pre-equilibrated with INSIDE buffer. For the assay, proteoliposomes were diluted into 120 mM NaCl, 5 mM HEPES (pH 6.8), 2 mM MgSO_4_ (OUTSIDE buffer). It was noted that 5-aminolevulinic acid appears to collapse the pH gradient across the liposome membranes.

A Cary Eclipse fluorescence spectrophotometer (Agilent Technologies) equipped with a cuvette and magnetic flea was used for the transport assays. Dual fluorescence excitation was set to 460/415 nm with emission at 510 nm. Transport was initiated following addition of 1 μM valinomycin. The resulting data was exported and analysed using Prism 7.0 (GraphPad Software).

### Competition assays using radiolabelled peptide.

Proteoliposomes were harvested as described above, the resulting pellet was resuspended in INSIDE buffer, subjected to three freeze-thaw cycles in liquid nitrogen, extruded through a 0.4 μm polycarbonate membrane and harvested by ultracentrifugation. For the assay, proteoliposomes were diluted in 110 mM NaCl, 10 mM NaPO_4_ pH 6.8 and 2 mM MgSO_4_ (OUTSIDE buffer) supplemented with 50 μM di-alanine containing trace amount of ^3^H di-alanine and the competing substrate at increasing concentrations. Competition assays were performed at 30 °C, with samples taken at specified time points, transport was halted by dilution into 2 mL cold H_2_O. Proteoliposomes were immediately filtered onto a 0.22 μm cellulose filter (Merck Millipore) using a vacuum manifold and subsequently washed twice more with 2 mL cold H_2_O. The amount of peptide transported into the liposomes was calculated based on the specific activity for each peptide using a Wallac scintillation counter. Experiments were performed a minimal of four times to generate an overall mean and S.D.

### Data availability

Atomic coordinates have been deposited in the Protein Data Bank (PDB) under accession numbers 6GZ9 (Valacyclovir complex), 6H7U and 6HZP (5-aminolevulinic acid complexes).

## Acknowledgements

We thank the staff of beamline ID23eh2 at the European Synchrotron Radiation Facililty (ESRF) and at beamline I24 Diamond Light Source, UK. This work was supported by Wellcome (102890/Z/13/Z).

## References

1. Giacomini KM, et al. (2010) Membrane transporters in drug development. Nat Rev Drug Discov 9(3):215-236.

2. Lin L, Yee SW, Kim RB, & Giacomini KM (2015) SLC transporters as therapeutic targets: emerging opportunities. Nat Rev Drug Discov 14(8):543-560.

3. Thomas VH, et al. (2006) The road map to oral bioavailability: an industrial perspective. Expert opinion on drug metabolism & toxicology 2(4):591-608.

4. Rubio-Aliaga I & Daniel H (2002) Mammalian peptide transporters as targets for drug delivery. Trends in pharmacological sciences 23(9):434-440.

5. Jornada DH, et al. (2015) The Prodrug Approach: A Successful Tool for Improving Drug Solubility. Molecules 21(1):42.

6. Rautio J, Meanwell NA, Di L, & Hageman MJ (2018) The expanding role of prodrugs in contemporary drug design and development. Nat Rev Drug Discov.

7. Brandsch M (2013) Drug transport via the intestinal peptide transporter PepT1. Current Opinion in Pharmacology 13(6):881-887.

8. Qumseya BJ, David W, & Wolfsen HC (2013) Photodynamic Therapy for Barrett’s Esophagus and Esophageal Carcinoma. Clin Endosc 46(1):30-37.

9. Anderson CMH, et al. (2010) Transport of the Photodynamic Therapy Agent 5‐ Aminolevulinic Acid by Distinct H+-Coupled Nutrient Carriers Coexpressed in the Small Intestine. Journal of Pharmacology and Experimental Therapeutics 332(1):220-228.

10. Sala-Rabanal M, Loo DDF, Hirayama BA, & Wright EM (2008) Molecular mechanism of dipeptide and drug transport by the human renal H+/oligopeptide cotransporter hPEPT2. American journal of physiology Renal physiology 294(6):F1422-1432.

11. Zaïr ZM, Eloranta JJ, Stieger B, & Kullak-Ublick GA (2008) Pharmacogenetics of OATP (SLC21/SLCO), OAT and OCT (SLC22) and PEPT (SLC15) transporters in the intestine, liver and kidney. Pharmacogenomics 9(5):597-624.

12. Brandsch M, Knütter I, & Bosse-Doenecke E (2008) Pharmaceutical and pharmacological importance of peptide transporters. Journal of Pharmacy and Pharmacology 60(5):543-585.

13. Terada T & Inui K-I (2004) Peptide transporters: structure, function, regulation and application for drug delivery. Current drug metabolism 5(1):85-94.

14. Perry CM & Faulds D (1996) Valaciclovir. A review of its antiviral activity, pharmacokinetic properties and therapeutic efficacy in herpesvirus infections. Drugs 52(5):754-772.

15. Yang B & Smith D (2012) Significance of Peptide Transporter 1 in the Intestinal Permeability of Valacyclovir in Wild-Type and PepT1 Knockout Mice. Drug metabolism and disposition: the biological fate of chemicals.

16. Sawada K, Terada T, Saito H, Hashimoto Y, & Inui KI (1999) Recognition of L-amino acid ester compounds by rat peptide transporters PEPT1 and PEPT2. The Journal of pharmacology and experimental therapeutics 291(2):705-709.

17. Ganapathy ME, Huang W, Wang H, Ganapathy V, & Leibach FH (1998) Valacyclovir: a substrate for the intestinal and renal peptide transporters PEPT1 and PEPT2. Biochemical and biophysical research communications 246(2):470-475.

18. Balimane PV, et al. (1998) Direct evidence for peptide transporter (PepT1)- mediated uptake of a nonpeptide prodrug, valacyclovir. Biochem Biophys Res Commun 250(2):246-251.

19. Incecayir T, et al. (2016) Carrier-Mediated Prodrug Uptake to Improve the Oral Bioavailability of Polar Drugs: An Application to an Oseltamivir Analogue. J Pharm Sci 105(2):925-934.

20. Varghese Gupta S, et al. (2011) Enhancing the Intestinal Membrane Permeability of Zanamivir: A Carrier Mediated Prodrug Approach. Molecular pharmaceutics 8(6):2358-2367.

21. Song X, et al. (2005) Amino acid ester prodrugs of the antiviral agent 2-bromo-5,6‐ dichloro-1-(beta-D-ribofuranosyl)benzimidazole as potential substrates of hPEPT1 transporter. Journal of medicinal chemistry 48(4):1274-1277.

22. Schlessinger A, et al. (2018) Molecular Modeling of Drug-Transporter Interactions-an International Transporter Consortium Perspective. Clin Pharmacol Ther.

23. Colas C, Ung PM, & Schlessinger A (2016) SLC Transporters: Structure, Function, and Drug Discovery. Medchemcomm 7(6):1069-1081.

24. Weitz D, et al. (2007) Functional and structural characterization of a prokaryotic peptide transporter with features similar to mammalian PEPT1. J. Biol. Chem. 282(5):2832-2839.

25. Newstead S (2015) Molecular insights into proton coupled peptide transport in the PTR family of oligopeptide transporters. Bba-Gen Subjects 1850(3):488-499.

26. Guettou F, et al. (2014) Selectivity mechanism of a bacterial homolog of the human drug-peptide transporters PepT1 and PepT2. Nature structural & molecular biology 21(8):728-731.

27. Daniel H, Spanier B, Kottra G, & Weitz D (2006) From bacteria to man: archaic proton-dependent peptide transporters at work. Physiology (Bethesda, Md) 21:93-102.

28. Steiner H-Y, Naider F, & Becker JM (1995) The PTR family: a new group of peptide transporters. Molecular microbiology 16(5):825-834.

29. Newstead S (2017) Recent advances in understanding proton coupled peptide transport via the POT family. Curr Opin Struct Biol 45:17-24.

30. Lyons JA, et al. (2014) Structural basis for polyspecificity in the POT family of proton-coupled oligopeptide transporters. EMBO Rep 15(8):886-893.

31. Martinez Molledo M, Quistgaard EM, Flayhan A, Pieprzyk J, & Low C (2018) Multispecific Substrate Recognition in a Proton-Dependent Oligopeptide Transporter. Structure 26(3):467-476 e464.

32. Parker JL, Mindell JA, & Newstead S (2014) Thermodynamic evidence for a dual transport mechanism in a POT peptide transporter. Elife 3.

33. Minhas GS, et al. (2018) Structural basis of malodour precursor transport in the human axilla. Elife 7.

34. Guo A, Hu P, Balimane PV, Leibach FH, & Sinko PJ (1999) Interactions of a nonpeptidic drug, valacyclovir, with the human intestinal peptide transporter (hPEPT1) expressed in a mammalian cell line. J Pharmacol Exp Ther 289(1):448-454.

35. Epling D, Hu Y, & Smith DE (2018) Evaluating the intestinal and oral absorption of the prodrug valacyclovir in wildtype and huPepT1 transgenic mice. Biochem Pharmacol 155:1-7.

36. Caffrey M (2011) Crystallizing membrane proteins for structure-function studies using lipidic mesophases. Biochemical Society transactions 39(3):725-732.

37. Solcan N, et al. (2012) Alternating access mechanism in the POT family of oligopeptide transporters. The EMBO journal 31(16):3411-3421.

38. Meredith D, et al. (2000) Modified amino acids and peptides as substrates for the intestinal peptide transporter PepT1. European journal of biochemistry / FEBS 267(12):3723-3728.

39. Pieri M, Gan C, Bailey P, & Meredith D (2009) The transmembrane tyrosines Y56, Y91 and Y167 play important roles in determining the affinity and transport rate of the rabbit proton-coupled peptide transporter PepT1. The International Journal of Biochemistry & Cell Biology 41(11):2204-2213.

40. Fowler PW, et al. (2015) Gating topology of the proton-coupled oligopeptide symporters. Structure 23(2):290-301.

41. Doki S, et al. (2013) Structural basis for dynamic mechanism of proton-coupled symport by the peptide transporter POT. Proc Natl Acad Sci U S A 110(28):11343-11348.

42. Bailey P, et al. (2000) How to Make Drugs Orally Active: A Substrate Template for Peptide Transporter PepT1. Angewandte Chemie (International ed in English) 39(3):505-508.

43. Lavis LD (2008) Ester bonds in prodrugs. ACS Chem Biol 3(4):203-206.

44. Aduri NG, et al. (2015) Salt Bridge Swapping in the EXXERFXYY Motif of Proton coupled Oligopeptide Transporters. The Journal of biological chemistry 290(50):29931-29940.

45. Samsudin F, Parker JL, Sansom MS, Newstead S, & Fowler PW (2016) Accurate Prediction of Ligand Affinities for a Proton-Dependent Oligopeptide Transporter. Cell Chem Biol 23(2):299-309.

46. Gardner LC & Cox TM (1988) Biosynthesis of heme in immature erythroid cells. The regulatory step for heme formation in the human erythron. The Journal of biological chemistry 263(14):6676-6682.

47. Stummer W, et al. (2006) Fluorescence-guided surgery with 5-aminolevulinic acid for resection of malignant glioma: a randomised controlled multicentre phase III trial. Lancet Oncol 7(5):392-401.

48. Neumann J & Brandsch M (2003) Delta-aminolevulinic acid transport in cancer cells of the human extrahepatic biliary duct. J Pharmacol Exp Ther 305(1):219-224.

49. Quistgaard EM, Martinez Molledo M, & Low C (2017) Structure determination of a major facilitator peptide transporter: Inward facing PepTSt from Streptococcus thermophilus crystallized in space group P3121. PLoS One 12(3):e0173126.

50. Parker JL, et al. (2017) Proton movement and coupling in the POT family of peptide transporters. Proc Natl Acad Sci U S A 114(50):13182-13187.

51. Newstead S, et al. (2011) Crystal structure of a prokaryotic homologue of the mammalian oligopeptide-proton symporters, PepT1 and PepT2. The EMBO journal 30(2):417-426.

52. Miroux B & Walker JE (1996) Over-production of proteins in Escherichia coli: mutant hosts that allow synthesis of some membrane proteins and globular proteins at high levels. Journal of Molecular Biology 260(3):289-298.

53. Caffrey M & Cherezov V (2009) Crystallizing membrane proteins using lipidic mesophases. Nature Protocols 4(5):706-731.

54. Kabsch W (2010) XDS. Acta Crystallographica Section D Biological Crystallography 66(Pt 2):125-132.

55. Evans PR & Murshudov GN (2013) How good are my data and what is the resolution? Acta Crystallographica Section D Biological Crystallography 69:1204-1214.

56. McCoy AJ, et al. (2007) Phaser crystallographic software. Journal of Applied Crystallography 40(Pt 4):658-674.

57. Emsley P, Lohkamp B, Scott WG, & Cowtan K (2010) Features and development of Coot. Acta Crystallographica Section D Biological Crystallography 66(Pt 4):486-501.

58. Adams PD, et al. (2010) PHENIX: a comprehensive Python-based system for macromolecular structure solution. Acta Crystallographica Section D Biological Crystallography 66(Pt 2):213-221.

59. Chen VB, et al. (2010) MolProbity: all-atom structure validation for macromolecular crystallography. Acta Crystallographica Section D Biological Crystallography 66(Pt 1):12-21.

